# Long-Term *In Vivo* Biocompatibility of Single-Walled Carbon Nanotubes

**DOI:** 10.1101/869750

**Authors:** Thomas V. Galassi, Merav Antman-Passig, Zvi Yaari, Jose Jessurun, Robert E. Schwartz, Daniel A. Heller

## Abstract

Over the past two decades, measurements of carbon nanotube toxicity and biodistribution have yielded a wide range of results. Properties such as nanotube type (single-walled vs. multi-walled), purity, length, aggregation state, and functionalization, as well as route of administration, greatly affect both the biocompatibility and biodistribution of carbon nanotubes. These differences suggest that generalizable conclusions may be elusive and that studies must be material- and application-specific. Here, we assess the short- and long-term biodistribution and biocompatibility of a single-chirality DNA-encapsulated single-walled carbon nanotube complex upon intravenous administration that was previously shown to function as an in-vivo reporter of endolysosomal lipid accumulation. Regarding biodistribution and fate, we found bulk specificity to the liver and >90% signal attenuation by 14 days in mice. Using near-infrared hyperspectral microscopy to measure single nanotubes, we found low-level, long-term persistence in organs such as the heart, liver, lung, kidney, and spleen. Measurements of histology, animal weight, complete blood count, and biomarkers of organ function all suggest short- and long-term biocompatibility. This work suggests that carbon nanotubes can be used as preclinical research tools in-vivo without affecting acute or long-term health.

## 1. Introduction

The unique physical properties of carbon nanotubes have prompted interest in many fields and potential applications in materials, electronics, biology, and medicine. Carbon nanotubes may exist as either single-walled carbon nanotubes (SWCNTs) or multi-walled carbon nanotubes (MWCNTs). SWCNTs are cylindrical graphene tubes with typical diameters of 0.5-2 nm and exhibit novel optical and electronic properties. These properties are dependent on the roll-up angle and diameter of nanotubes, which are defined by the integers (*n,m*), denoting SWCNT species/chirality^1^. MWCNTs consist of multiple concentric layers of graphene cylinders, and also exist as multiple chiralities exhibiting unique properties, with diameters that can range from 1-60 nm^2, 3^.

The unique properties of carbon nanotubes and their potential applications has prompted many studies of the toxicity and biological/ecological fate of these materials over the past two decades^4, 5^ that has often resulted in perceived general toxicity by many in the scientific community. Overall, however, the results of toxicology studies on carbon nanotubes have been remarkably inconsistent^4^. One major conflating factor in determining the biodistribution and toxicological profile of a nanotube sample is the type of nanotube under investigation. Studies investigating the effects of MWCNTs, for instance, suggest that they cause asbestos-like effects in mammals^6^, while recent investigations of SWCNTs found a protective effect against several disorders, including neurodegenerative disease and stroke^7–9^. Studies also found differences between SWCNTs and MWCNTs in both cell uptake and viability, with MWCNT preparations leading to a decrease in cell viability that was not seen upon exposure with SWCNT preparations^10^. Further investigation into these results reveals that there are many factors related to carbon nanotube structure, size, preparation techniques, and route of administration that affect the biodistribution and toxicological profile of a specific carbon nanotube preparation^7–9^.

Studies focusing on SWCNTs report a large variety of findings ranging from harmful to beneficial effects on cells or animals, depending on the preparation/derivatization. Recent work investigating the structure-dependent biocompatibility of SWCNTs found that, while singly-dispersed SWCNTs showed no significant effects on mitochondrial function and hypoxia *in vitro*, negative effects were seen upon treatment with aggregated SWCNTs which further depended on the structural integrity of the aggregated SWCNT dispersions^11^. Aggregated SWCNTs were also found, however, to attenuate the effects of methamphetamine in mice^8^. Recent work also suggests that SWCNT treatment alleviates autophagic/lysosomal defects in primary glia from a mouse model of Alzheimer’s disease^7^, while amine-modified SWCNTs are neuroprotective in a stroke model in rats^9^. Individually-dispersed SWCNTs functionalized with lipid-polyethylene glycol (PEG) conjugates, injected intravenously into rodents, exhibited minimal effects on rodent blood chemistry^12^, despite their presence in the liver for up to four months. Such studies also highlight the effects of SWCNT functionalization on long term biodistribution. While the non-covalently functionalized SWCNTs persisted in organs such as the liver and spleen for months^12^, covalently functionalized SWCNTs were shown to rapidly clear the body via the urine^13^. Investigators have studied SWCNT toxicity in animals following airway, intravenous, intraperitoneal, subcutaneous, oral, and topical exposure, leading to varying degrees of biocompatibility depending on the SWCNT preparation used^5^. Even when focusing on a single route of administration, such as intravenous injection, it has been difficult to draw general conclusions about SWCNT toxicity due to large differences in the quantity of SWCNTs injected as well as varying preparation techniques. Injected quantities of SWCNTs have ranged over three orders of magnitude. The large range of concentrations and the diverse functionalization chemistries likely contribute to large differences in biodistribution and biocompatibility^13–17^

The diversity of results pertaining to carbon nanotube biodistribution and toxicity highlights the need for specific studies for each type, functionalization, and application of carbon nanotubes. Here, we examine the short and long-term biodistribution and toxicity of a single, purified SWCNT (n,m) species/chirality (the (9,4) species) encapsulated with a specific single-stranded DNA sequence (ssCTTC_3_TTC; DNA-SWCNT). This DNA-nanotube combination, when injected intravenously into mice, was recently found to function as a non-invasive reporter of Kupffer cell endolysosomal lipid accumulation^17^. Bulk measurements have shown that this DNA-nanotube complex, ssCTTC_3_TTC-(9,4), localized predominantly to the liver, although single-particle measurements found small quantities in other organs. Extensive histological examinations, complete blood counts, and serum biomarker measurements results suggest that ssCTTC_3_TTC-(9,4) is sufficiently biocompatible for preclinical applications, while more work is needed to assess its potential for use in humans.

## 2. Experimental Section

### 2.1 Preparation of the purified DNA-nanotube complex, ssCTTC_3_TTC-(9,4)

1 mg/mL of raw EG 150X single walled carbon nanotubes purchased from Chasm Advanced Materials (Norman, Oklahoma) were mixed with 2 mg/mL ssDNA in 0.1 M NaCl. SWCNTs were then wrapped in ssDNA via probe tip ultrasonication (Sonics & Materials, Inc.) for 120 minutes at ~ 8 W. Dispersions were then centrifuged (Eppendorf 5430 R) for 90 minutes at 17,000 x g. The top 85% of the resulting supernatant was then collected and used for purification of the (9,4) chirality.

Purification of the (9,4) nanotube from the unsorted ssCTTC_3_TTC-SWCNT sample was performed using the aqueous two-phase extraction method^18–20^. In brief, ssCTTC_3_TTC-SWCNT was mixed with a solution containing a final concentration of 7.76% polyethylene glycol (PEG, molecular weight 6 kDa, Alfa Aesar), and 15.0% polyacrylamide (PAM, molecular weight 10 kDa, Sigma Aldrich). Following an overnight incubation at room temperature, the sample was vortexed and then centrifuged at 10,000 x g for 3 minutes. The top phase of the resulting solution was then collected and added to blank “bottom phase,” which was produced by removing the bottom phase of a 7.76 PEG, 15.0% PAM solution following centrifugation at 10,000 x g for 10 minutes. The resulting solution was once again vortexed and centrifuged to produce a top phase enriched in ssCTTC_3_TTC-(9,4) complexes. Following collection of the top phase NaSCN was added at a final concentration of 0.5 M. This solution was then incubated overnight at 4 degrees Celsius to precipitate ssCTTC_3_TTC-(9,4). The sample was then centrifuged at 17,000 x g for 20 minutes, causing ssCTTC_3_TTC-(9,4) to pellet. Following removal of the supernatant, this pellet was then suspended in diH_2_O and stored with 0.1 mg/mL free ssCTTC_3_TTC-(9,4).

### 2.2 Animal Studies

All animal studies were approved and carried out in accordance with the Memorial Sloan Kettering Cancer Center Institutional Animal Care and Use Committee. All animals used in the study were male C57BL/6 mice at 6-12 weeks of age. All control and experimental mice were age matched and housed in identical environments. For the assessment of ssCTTC_3_TTC-(9,4) *in vivo*, mice were tail vein injected with 200 μL of 0.5mg/L ssCTTC_3_TTC-(9,4) diluted in PBS. For all other experiments, mice were tail vein injected with 200 μL of 1.0 mg/L ssCTTC_3_TTC-(9,4) diluted in PBS. For *in vivo* spectroscopy, mice were anesthetized with 2% isoflurane prior to data collection.

### 2.3 Near infrared in vivo spectroscopy

Spectra were acquired from rodents non-invasively *in vivo* using a custom-built reflectance probe-based spectroscopy system^17, 21, 22^. The excitation was provided by injection of a 730 nm diode laser (Frankfurt) into a bifurcated fiber optic reflection probe bundle (Thorlabs). The sample leg of the bundle included one 200 μm, 0.22 NA fiber optic cable for sample excitation located in the center of six 200 μm, 0.22 NA fiber optic cables for collection of the emitted light. The exposure time for all acquired data was 5 seconds. Light below 1050 nm was removed via long pass filters, and the emission was focused through a 410 μm slit into a Czerny-Turner spectrograph with 303 mm focal length (Shamrock 303i, Andor). The beam was dispersed by an 85 g/mm grating with 1350 nm blaze wavelength and collected by an iDus InGaAs camera (Andor). After acquisition, data was processed to apply spectral corrections for non-linearity of the InGaAs detector response, background subtraction, and baseline subtraction *via* the use of OriginPro 9 software with a standard adjacent averaging smoothing method and a spline interpolation method.

### 2.4 Tissue fixation and sectioning

Organs were fixed in 10% buffered formalin phosphate and paraffin embedded. 5 μm sections were then placed on glass slides. Paraffin was removed and sections were either left unstained for near-infrared hyperspectral microscopy or stained with haematoxylin and eosin (H&E) for basic histology at the Molecular Cytology Core Facility of Memorial Sloan Kettering Cancer Center and Histowiz Inc. (Brooklyn, NY).

### 2.5 Near-Infrared hyperspectral microscopy

Near-infrared hyperspectral microscopy was performed as previously described^23^. In brief, a continuous wave 730 nm diode laser (output power = 2 W) was injected into a multimode fiber to provide an excitation source. After passing through a beam shaping optics module to produce a top hat intensity profile (maximum 20% variation on the surface of the sample), the laser was reflected through an inverted microscope (with internal optics modified for near-infrared transmission) equipped with a 100X (UAPON100XOTIRF, NA=1.49) oil objective (Olympus, USA) *via* a longpass dichroic mirror with a cut-on wavelength of 880 nm. Spatially resolved near-infrared emission was then passed twice through a turret-mounted volume Bragg grating (VBG) which allowed the light to be spectrally resolved. The monochromatic beam with 3.7 nm FWHM was collected by a 256 × 320 pixel InGaAs camera (Photon Etc) to result in an image. Spectrally-defined images were collected with a 4 s integration time. The VBG was rotated in 4 nm steps between 1100-1200 nm (26 images in total). Data rectification was conducted using PhySpec software (Photon Etc) to result in “hyperspectral cubes” wherein every pixel of a near-infrared image was spectrally resolved^23^.

### 2.6 Analysis and processing of hyperspectral data

Hyperspectral data was saved as 16 bit arrays (320 × 256 × 26) where the first two coordinates represent the spatial location of a pixel and the last coordinate its position in wavelength space. Background subtraction and intensity corrections to compensate for non-uniform excitation were applied via MATLAB code developed in the authors’ lab. ROIs were then manually selected from images and spectra were obtained using the Time Series Analyzer plugin for ImageJ.

## 3. Results

### 3.1 In vivo detection of ssCTTC_3_TTC-(9,4) DNA-nanotube complexes

The near-infrared signal from the purified DNA-nanotube complex was assessed transiently. Previous work describes the isolation of the DNA-nanotube complex, ssCTTC_3_TTC-(9,4), consisting of the single-stranded DNA sequence, CTTC_3_TTC, encapsulating the nanotube species (9,4), by aqueous two-phase extraction^20^. The complexes were found to localize to the liver of mice following intravenous injection, where it was non-invasively assessed via a fiber optic probe^17^. Spectra of the nanotube photoluminescence in mice were acquired using a fiber optic probe-coupled spectrometer and near-infrared camera (described in Methods). The measurements found a sharp decrease in nantoube fluorescence intensity over two weeks after injection (Figure 1). This result suggests that nanotubes were either quenched or excreted from the liver over this time period.

**Figure 1.**
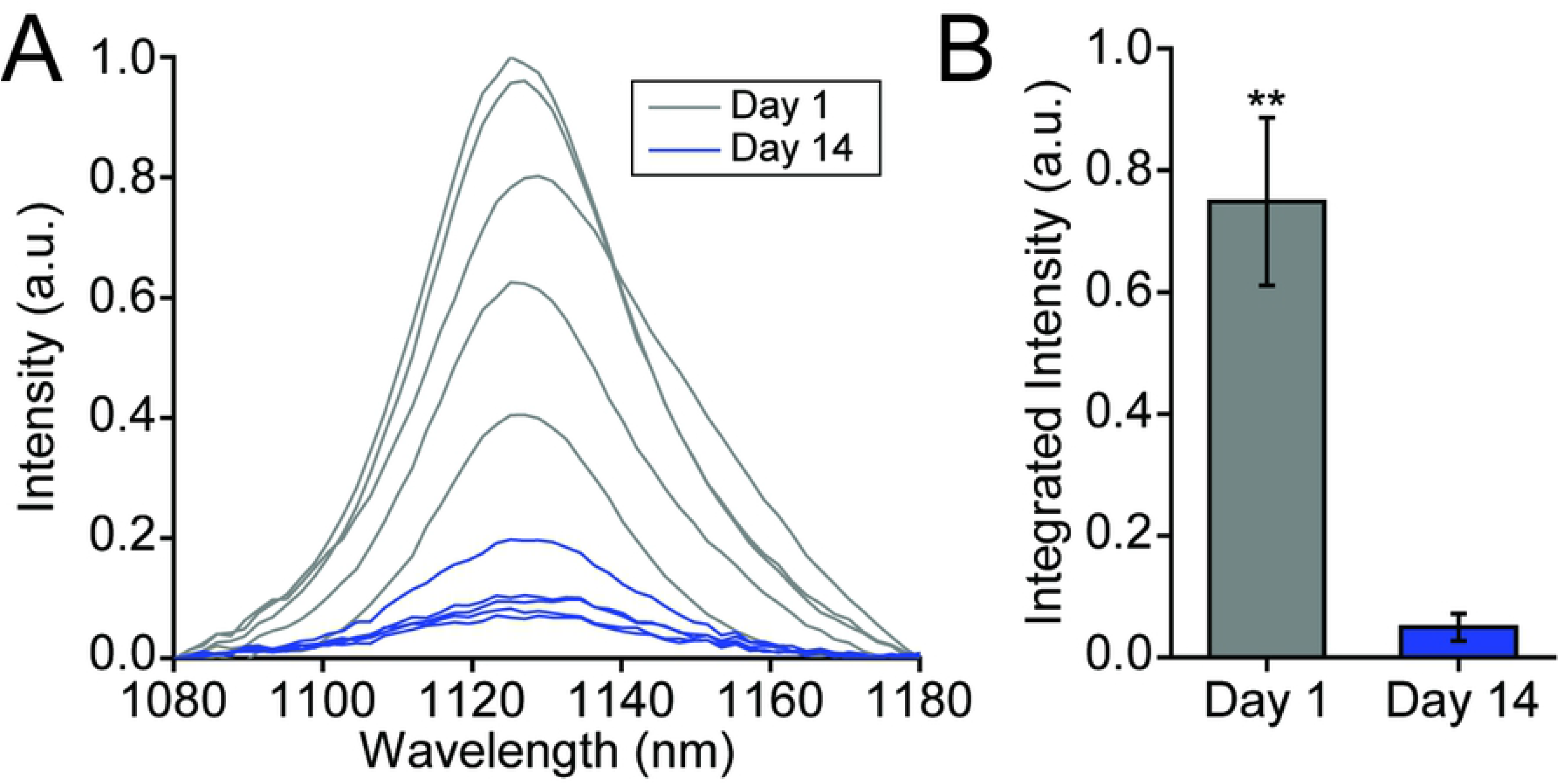
Near-infrared spectroscopy of single-species DNA-nanotube complexes *in vivo*. **A)** Near-infrared emission spectra of ssCTTC_3_TTC-(9,4) DNA-SWCNT complexes measured from the region of the mouse liver *in vivo* using a fiber optic probe device following intravenous injection into a mouse. **B)** Normalized integrated intensity of spectra depicted in (A). Error bars are standard deviation from N=5 mice. **=P<.01 as measured with a Student’s t-test.

### 3.2 Microscopy and histology of SWCNTs in resected murine tissues

To further investigate the biodistribution of ssCTTC_3_TTC-(9,4) DNA-nanotube complexes following intravenous injection, we examined tissue sections via near-infrared hyperspectral microscopy. This technique can image single SWCNTs^24^ in tissues even in the presence of normal tissue autofluorescence^25–27^. We resected and paraffinized tissue sections of heart, liver, lung, kidney, spleen and brain 24 hours after administration of SWCNTs and imaged using the near-infrared hyperspectral microscope at 100X magnification. Upon investigation of the near-infrared data, individual ROIs, denoting ssCTTC_3_TTC-(9,4) complexes, could be detected in liver, spleen, lung, kidney and heart, with highest prevalence in liver and spleen (Figure 2). The same tissues were also processed via H&E staining (Figure 2). The stained tissues were assessed by a trained pathologist; no signs of tissue injury or other abnormalities were found. No signal was detected in the brain via *in vivo* measurements (Figure S1) or in tissue sections (Figure 2).

**Figure 2.**
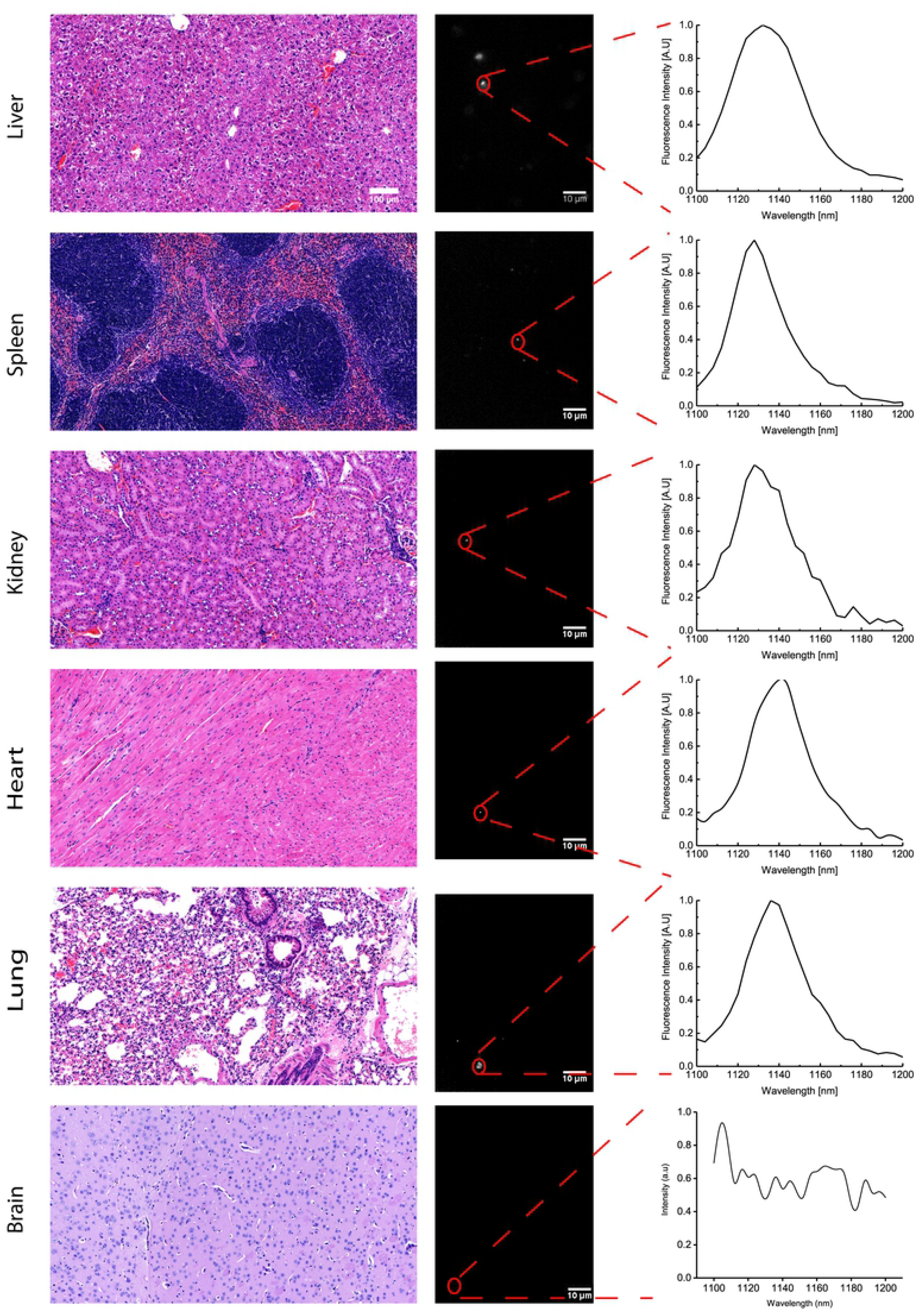
Imaging carbon nanotubes in murine tissues 24 hours after injection. H&E stains (left) and hyperspectral microscopy images (middle) of various organs 24 hours after intravenous injection with ssCTTC_3_TTC-(9,4) complexes. Representative fluorescence spectra (right) of the denoted complexes are shown.

We also assessed the long-term biodistribution of the nanotubes. Previous studies found the persistence of non-covalently functionalized SWCNTs in organs, such as the liver and spleen, for several months^16^. We used near-infrared hyperspectral microscopy to assess the long term biodistribution of ssCTTC_3_TTC-(9,4) in heart, liver, lung, kidney, and spleen tissue. One month following injection, nanotube emission could be seen in all tissue sections (Figure 3). The nanotubes were sparsely distributed through lung, heart, and kidney tissue, and more prevalent in the liver and spleen. Upon observation of the H&E stained tissue sections (Figure 3), no abnormalities were noted by a trained pathologist.

**Figure 3.**
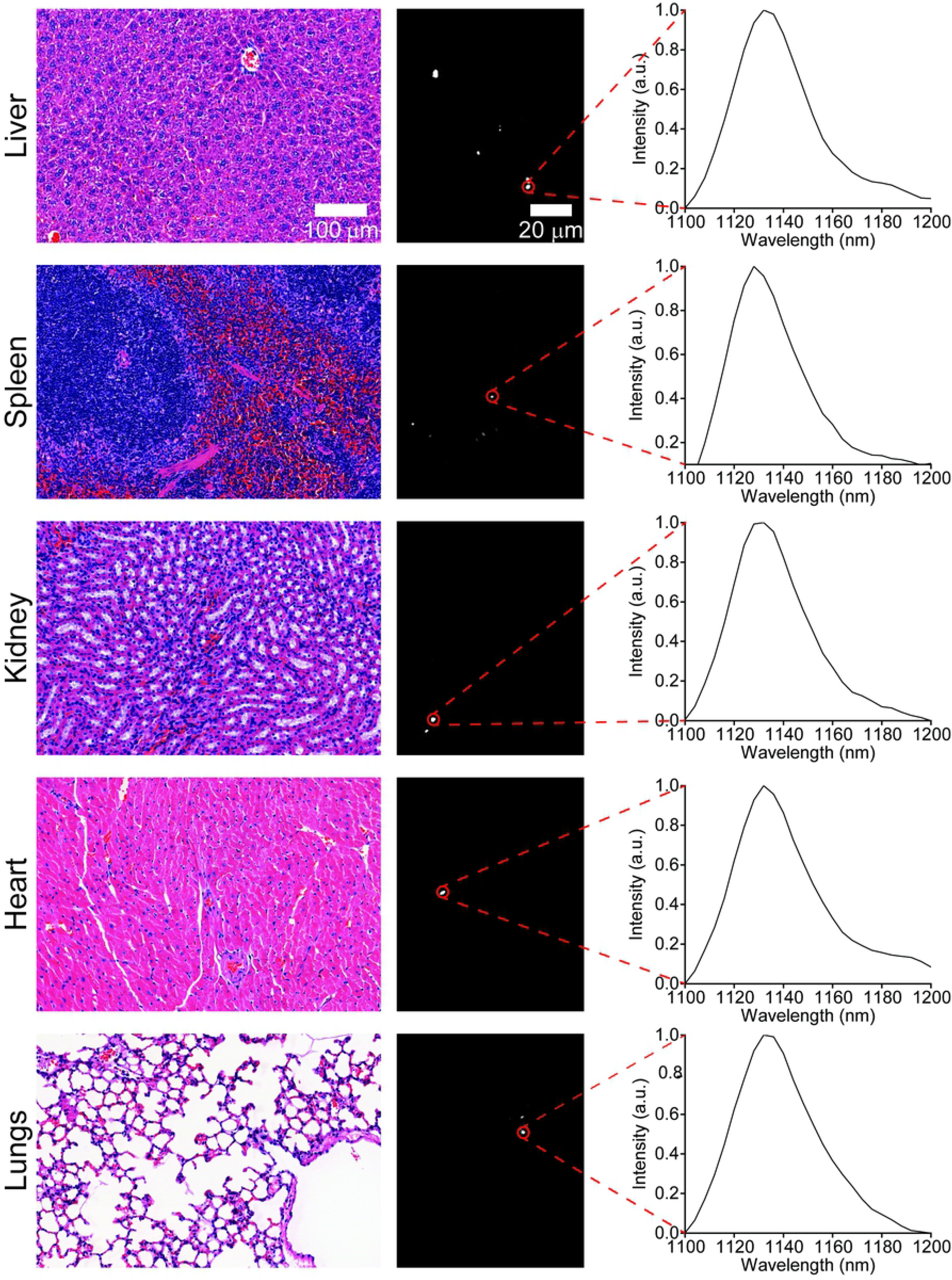
Imaging carbon nanotubes in murine tissues one month after injection. H&E stains (left) and hyperspectral microscopy images (middle) of various organs one month after intravenous injection with ssCTTC_3_TTC-(9,4) complexes. Representative fluorescence spectra (right) of the denoted complexes are shown.

Murine tissues were also assessed at three and five months after injection. No nanotubes were found in lung tissue at the three-month timepoint (Figure 4), or in lung or heart tissue at five months (Figure 5), although they were found in the liver, spleen, and kidneys at both timepoints. Despite chronic exposure to ssCTTC_3_TTC-(9,4) in these organs, no tissue abnormalities were observed upon histological inspection of these tissues by a trained pathologist (Figures 4-5).

**Figure 4.**
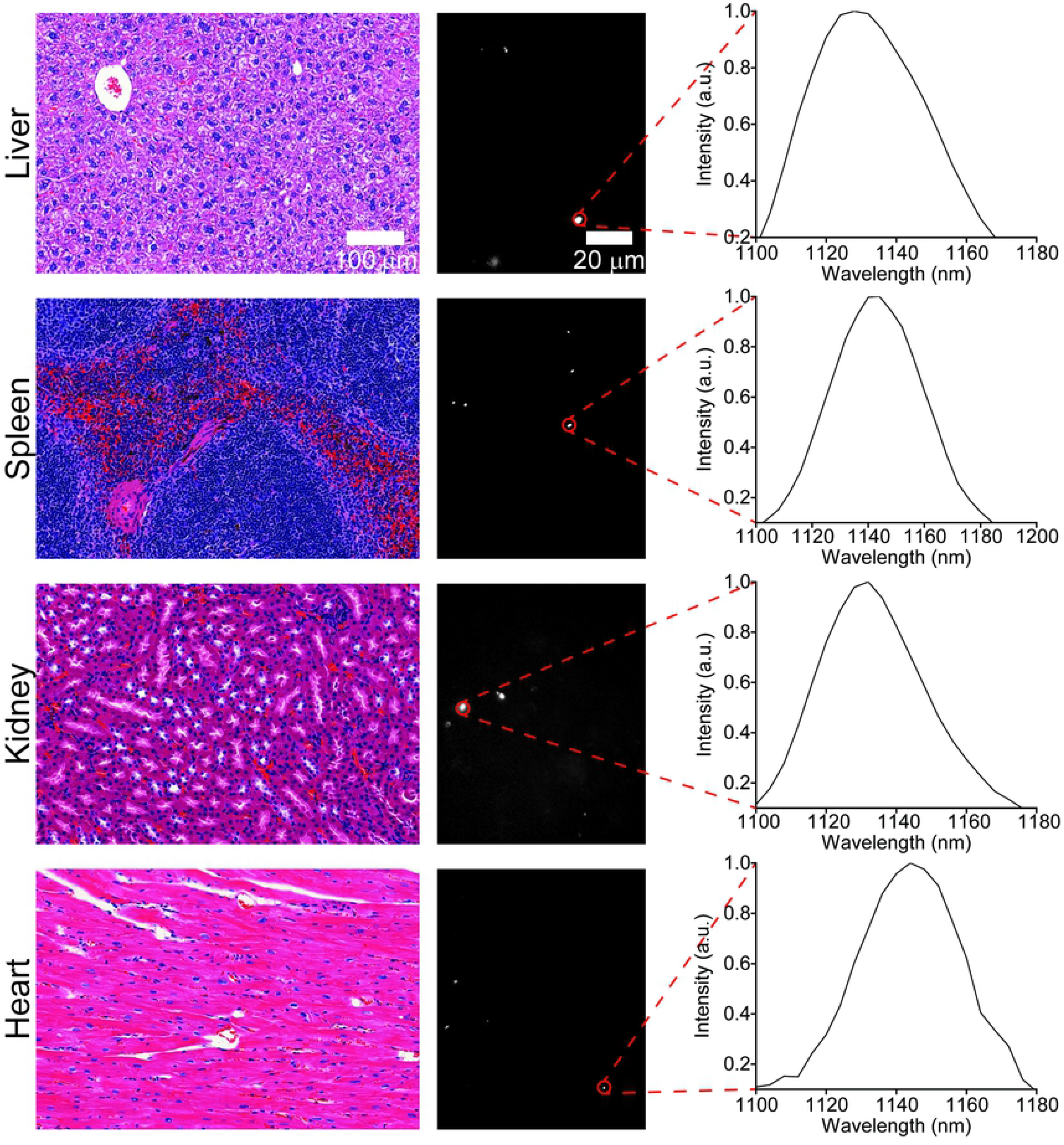
Imaging carbon nanotubes in murine tissues three months after injection. H&E stains (left) and hyperspectral microscopy images (middle) of various organs one month after intravenous injection with ssCTTC_3_TTC-(9,4) complexes. Representative fluorescence spectra (right) of the denoted complexes are shown.

**Figure 5.**
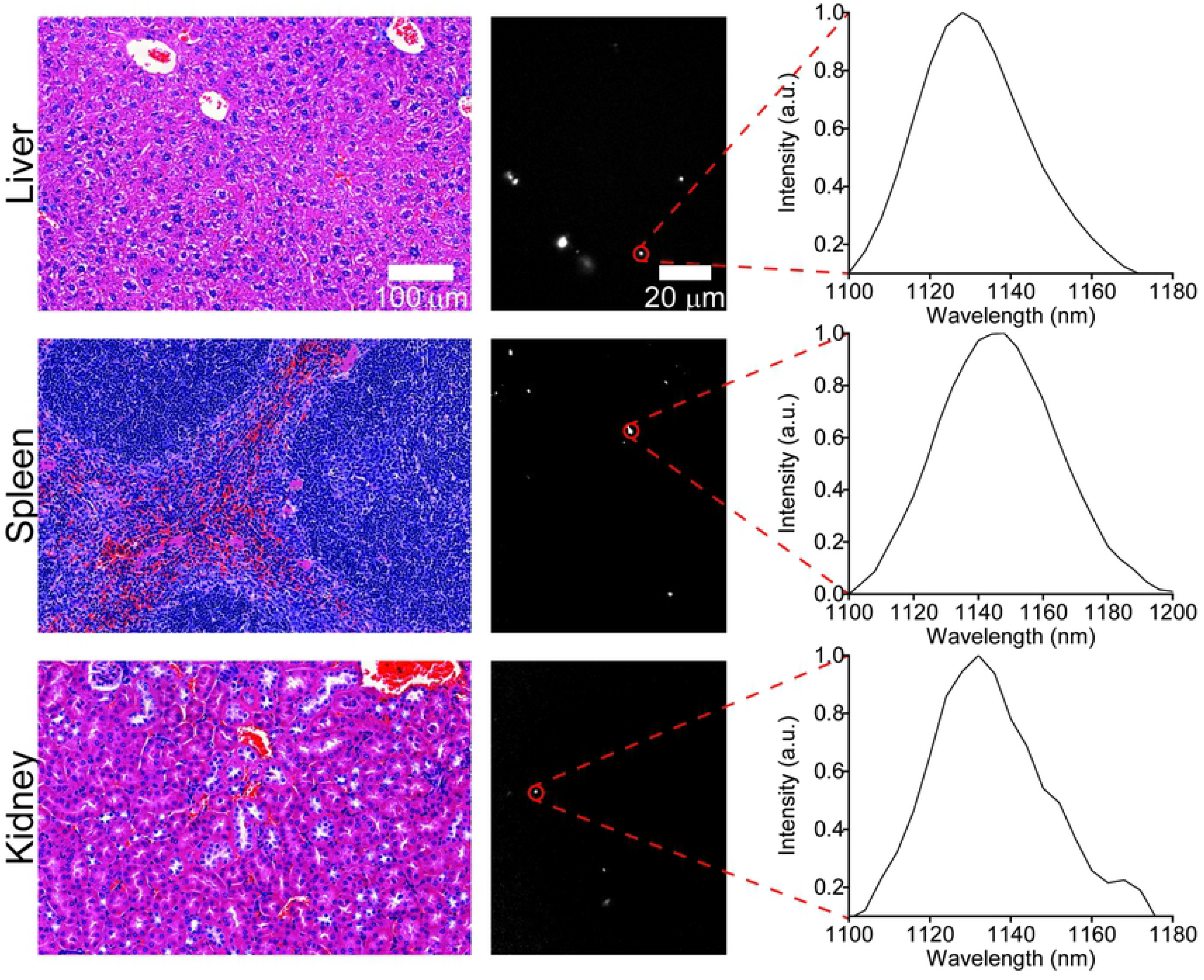
Imaging carbon nanotubes in murine tissues five months after injection. H&E stains (left) and hyperspectral microscopy images (middle) of various organs one month after intravenous injection with ssCTTC_3_TTC-(9,4) complexes. Representative fluorescence spectra (right) of the denoted complexes are shown.

**Figure 6.**
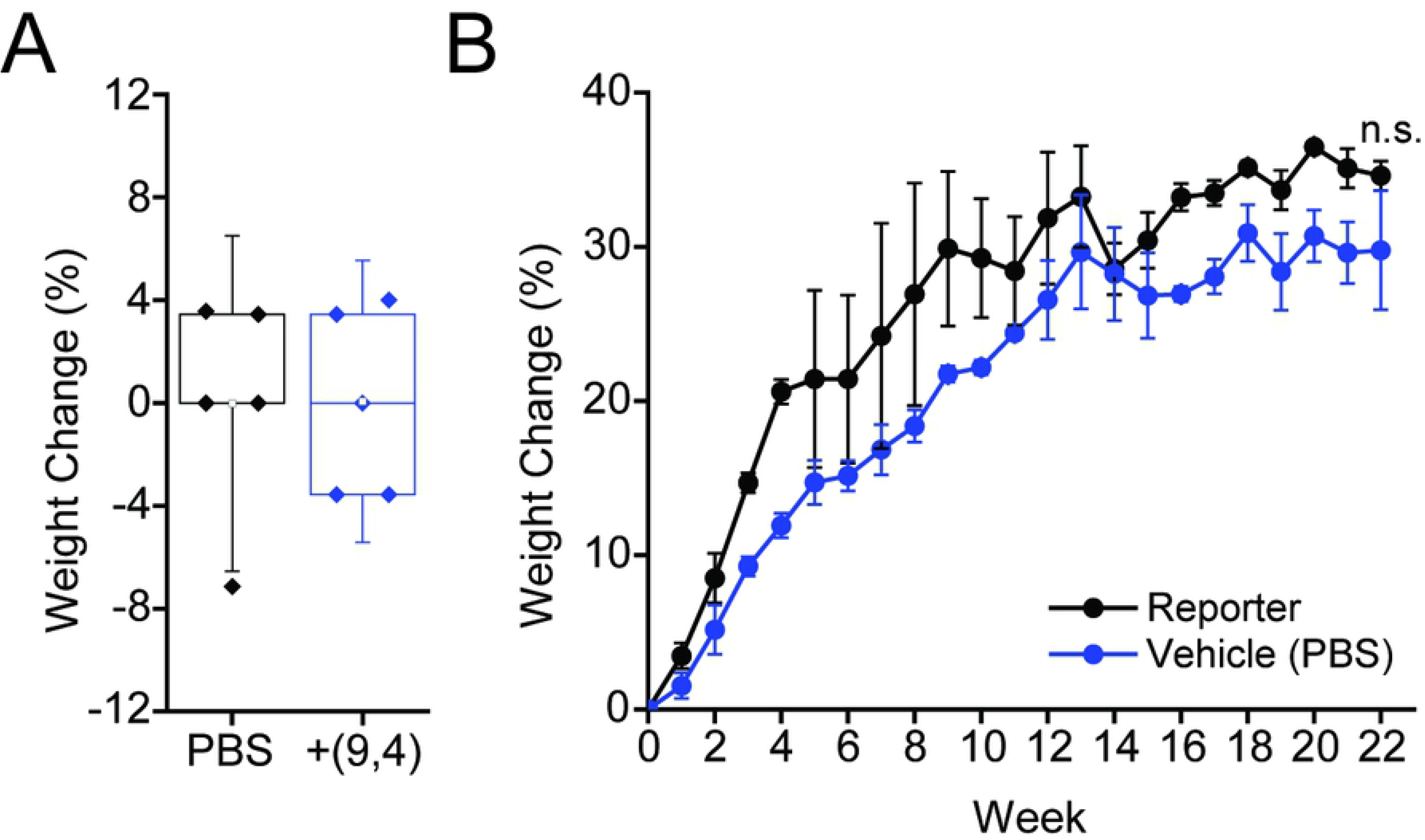
Effects of nanotubes on mouse weight. **A)** Weight change in mice 24 hours after injection with ssCTTC_3_TTC-(9,4) complexes or vehicle control (PBS). **B)** Weight change in mice followed 22 weeks after injection with ssCTTC_3_TTC-(9,4) complexes or vehicle control (PBS).

### 3.3 Mouse weight measurements

Mouse weights were measured after injection of the nanotube complexes over a period of five months. No significant difference was seen in weight changes 24 hours after injection with ssCTTC_3_TTC-(9,4) complexes (Figure 5A), consistent with previous results^17^. Similarly, mouse growth was not affected over a period of five months (Figure 5B).

### 3.4 Serum biomarker assessments

To further assess the effects of short and long-term exposure of the DNA-nanotube complexes, serum biomarkers, and complete blood counts were measured 24 hours and five months after administration of the nanotubes. Biomarkers of hepatic injury were measured 24 hours and 5 months after injection of ssCTTC_3_TTC-(9,4). Between nanotube and PBS injected mice, no statistically significant differences were found for the biomarkers alanine aminotransferase (ALT), aspartate aminotransferase (AST), globulin (GLOB), albumin (ALB), total protein, or TCO_2_ (Figure S1). A small, statistically significant increase was apparent in serum levels of serum alkaline phosphatase (Figure 7). However this slight difference is likely not biologically significant, as the values seen for nanotube injected mice here are consistent with those seen in another study that assessed ALP levels in similarly aged, male C57BL/6 mice injected with PBS^17^. A statistically significant difference is also apparent in the ALB:GLOB ratio in serum; this difference is slight and likely due to instrument limitations, as inspection of the raw data reveals that limited resolution in ALB:GLOB measurements is likely the cause for this significant difference (Figure 7). Despite the presence of nanotubes in the liver 5 months after injection, no signs of liver injury were evident from hepatic biomarkers (Figure 8).

**Figure 7.**
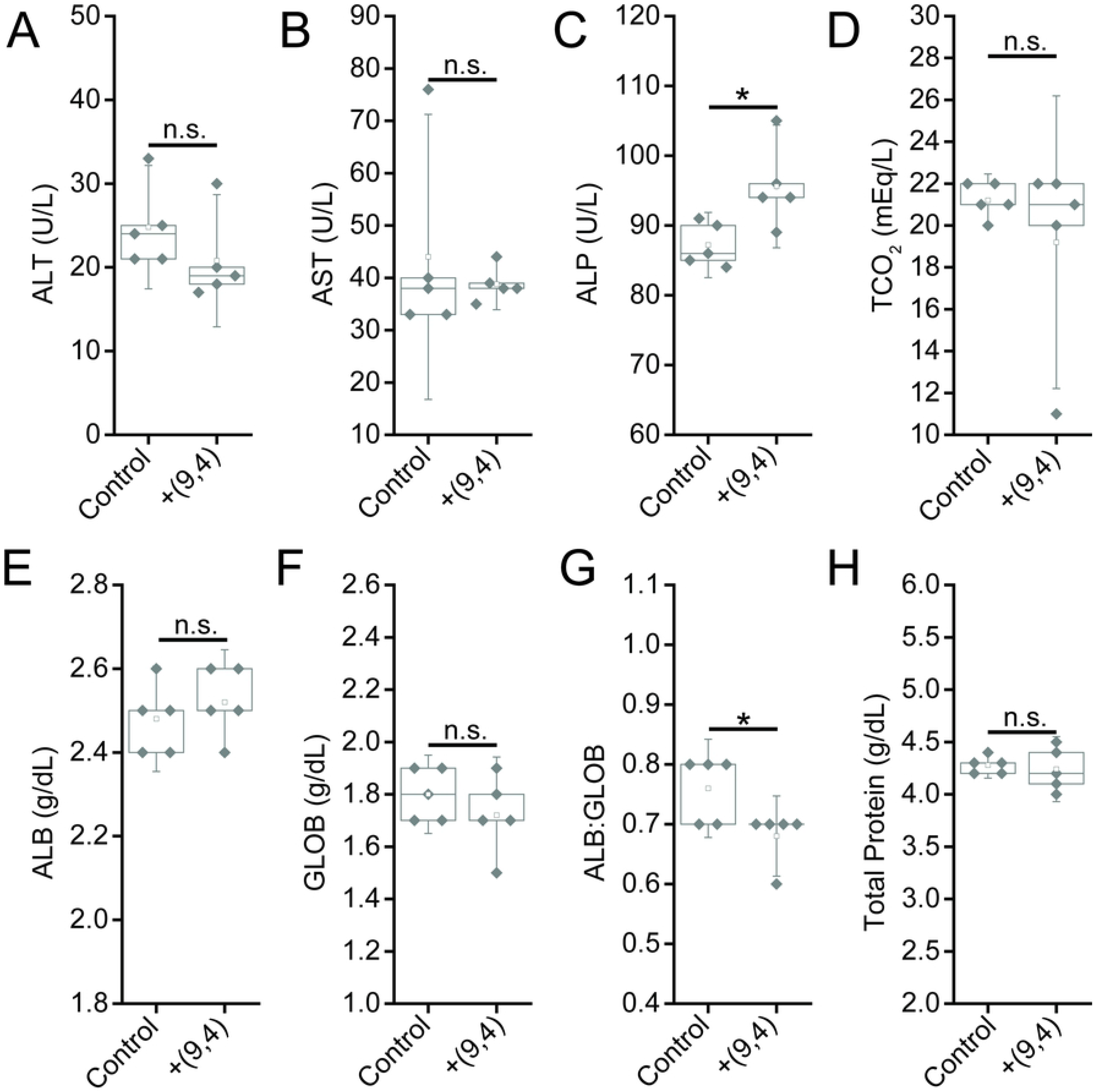
Serum chemistry measurements of biomarkers of hepatic injury in mice 24 hours after injection of nanotubes. Samples were measured 24 hours after injection of PBS (control) or ssCTTC_3_TTC-(9,4) DNA-nanotube complexes. **A)** Serum alanine transaminase concentrations (ALT) in mice. **B)** Serum aspartate transaminase (AST) concentrations in mice. **C)** Serum alkaline phosphatase (ALP) concentrations in mice. **D)** Serum carbon dioxide (TCO_2_) levels in mice. **E)** Serum albumin (ALB) levels in mice. **F)** Serum globulin (GLOB) levels in mice. **G)** Serum ALB:GLOB ratio in mice. **H)** Serum total protein levels in mice. * = p < .05 as determined with a Student’s two way t-test. N=5 mice per group.

**Figure 8.**
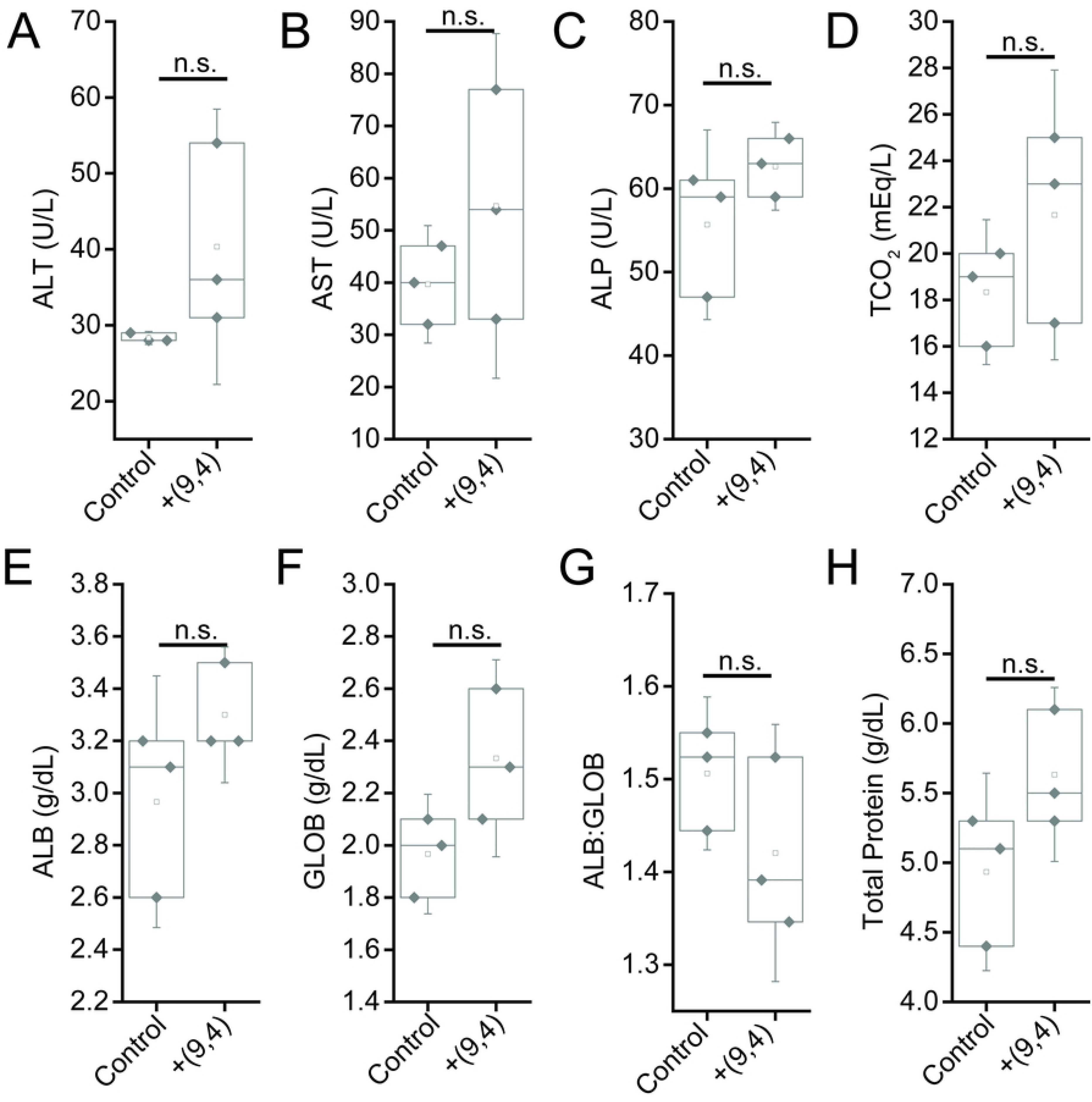
Serum chemistry measurements of biomarkers of hepatic injury in mice 5 months after injection of nanotubes. Samples were measured 5 months after injection of PBS (control) or ssCTTC_3_TTC-(9,4) DNA-nanotube complexes. **A)** Serum alanine transaminase concentrations (ALT) in mice. **B)** Serum aspartate transaminase (AST) concentrations in mice. **C)** Serum alkaline phosphatase (ALP) concentrations in mice. **D)** Serum carbon dioxide (TCO_2_) levels in mice. **E)** Serum albumin (ALB) levels in mice. **F)** Serum globulin (GLOB) levels in mice. **G)** Serum ALB:GLOB ratio in mice. **H)** Serum total protein levels in mice. Statistical significance was determined with a Student’s two way t-test. N=3 mice per group.

Serum biomarkers of renal function were assessed 24 hours and five months after injection of the nanotube complexes. Between nanotube-injected and control mice after 24 hours, no statistically significant differences were found for the biomarkers blood urea nitrogen (BUN), creatinine (CREA), BUN:CREA, phosphate (P), sodium (Na), Na:K, and anion gap (Figure 9). While chloride levels were raised in nanotube-injected mice, the increase was less than 2% (Figure 9). Potassium (K) levels were also different between the two groups, although they were actually lower in nanotube injected mice (Figure 9) which is not a sign of renal injury (an increase in K levels would be). Finally, a significant difference was seen in the Na:K ratio between groups. A decrease in this ratio is indicative of liver/kidney stress, however, rather than the slight increase seen herein (Figure 9). Overall, biomarkers of renal function at 24 hours suggest that the nanotubes did not cause any injury. After five months, Both BUN, CREA, and measured anion gaps were slightly increased five months after injection, with other renal biomarkers showing consistent, albeit slight (and statistically non-significant) changes (Figure 10), suggesting that future work is needed to assess the effects of the long-term persistence of ssCTTC_3_TTC-(9,4) on renal function.

**Figure 9.**
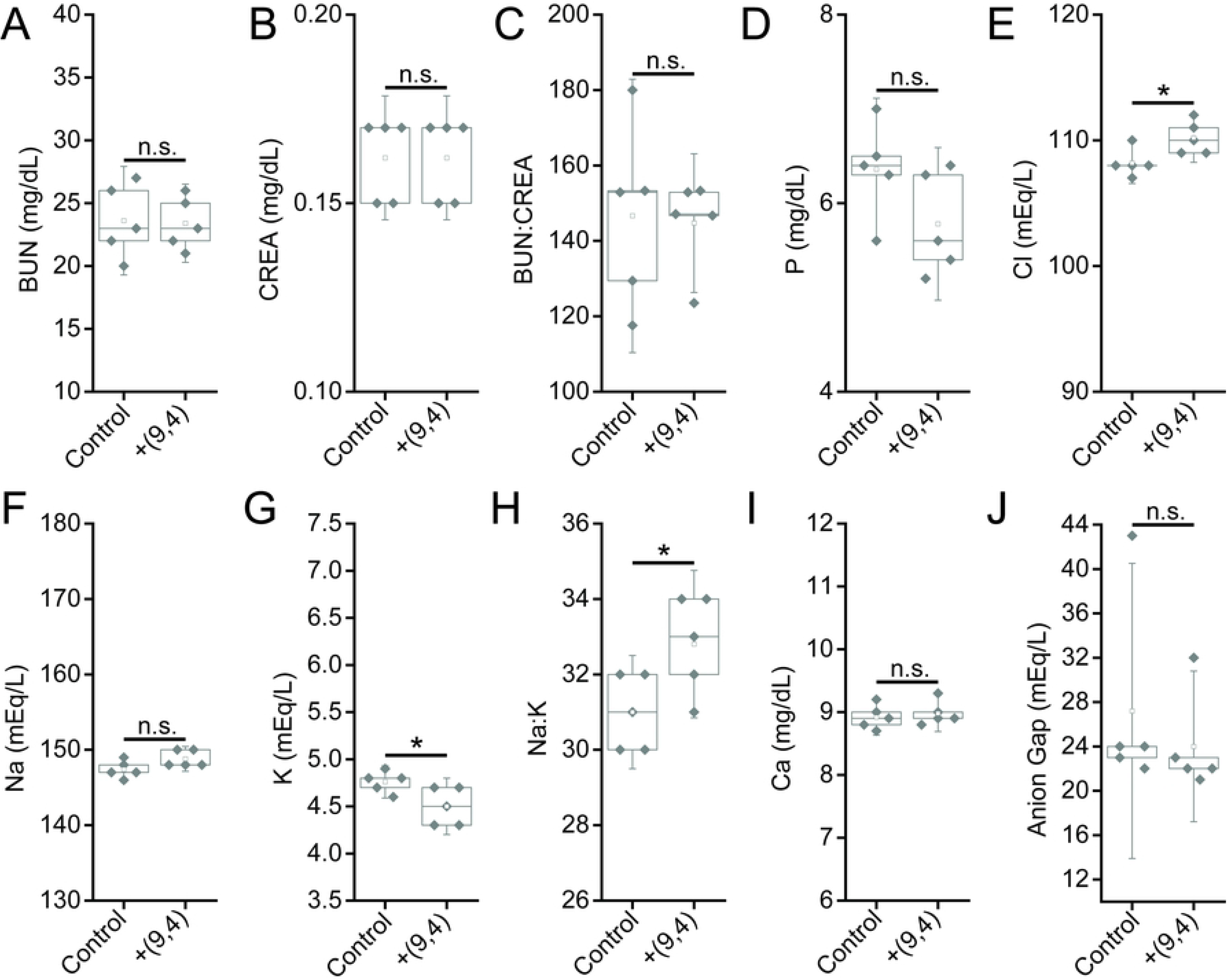
Serum chemistry measurements of biomarkers of renal function in mice 24 hours after injection of nanotubes. Samples were measured 24 hours after injection of PBS (control) or ssCTTC_3_TTC-(9,4) DNA-nanotube complexes. **A)** Blood urea nitrogen (BUN) concentration in mice. **B)** Serum creatinine (CREA) concentrations in mice. **C)** Serum BUN:CREA ratios in mice. **D)** Serum phosphate (P) concentration in mice. **E)** Serum chloride (Cl) concentrations in mice. **F)** Serum sodium (Na) concentrations in mice. **G)** Serum potassium (K) concentration in mice. **H)** Serum Na:K ratio in mice. **I)** Serum calcium (Ca) concentration in mice. **J)** Serum anion gap in mice. * = p < .05 as determined with a Student’s two way t-test. N=5 mice per group.

**Figure 10.**
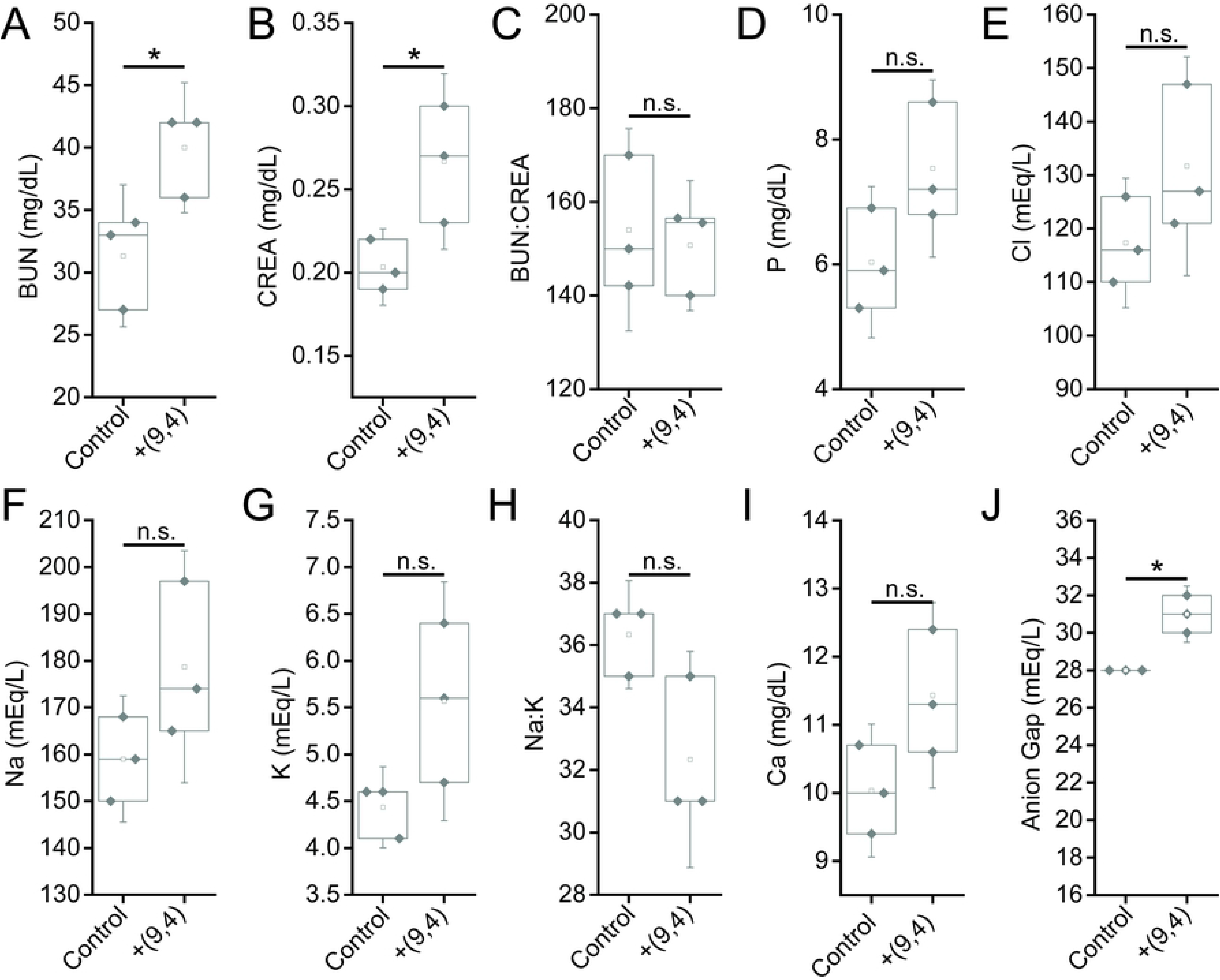
Serum chemistry measurements of biomarkers of renal function in mice 5 months after injection of nanotubes. Samples were measured 5 months after injection of PBS (control) or ssCTTC_3_TTC-(9,4) DNA-nanotube complexes. **A)** Blood urea nitrogen (BUN) concentration in mice. **B)** Serum creatinine (CREA) concentrations in mice. **C)** Serum BUN:CREA ratios in mice. **D)** Serum phosphate (P) concentration in mice. **E)** Serum chloride (Cl) concentrations in mice. **F)** Serum sodium (Na) concentrations in mice. **G)** Serum potassium (K) concentration in mice. **H)** Serum Na:K ratio in mice. **I)** Serum calcium (Ca) concentration in mice. **J)** Serum anioin gap in mice. * = p < .05 as determined with a Student’s two way t-test. N=3 mice per group.

Complete blood counts (CBCs) were performed 24 hours and 5 months after injection with the nanotube complexes. Counts of white blood cells (WBCs), lymphocytes, eosinophils, and basophils did not vary significantly, while there was a slight decrease in neutrophil count 24 h after injection (Figure 11). The number of monocytes increased slightly after 24 h, however the lack of a corresponding increase in neutrophils and other inflammatory markers coupled with the normal tissue architecture observed suggests that this slight difference is the result of normal biological variation and not likely due to the injection of nanotubes (Figures 2, 11). Blood counts associated with oxygen levels appeared normal; significant differences were not seen in red blood cell (RBC) counts, hemoglobin levels, hematocrit percentage, mean corpuscular hemoglobin quantity or concentration, or RBC distribution width (Figure 12). A slight, statically significant decrease was seen in platelet count between groups. This difference was not apparent at 5 months after administration (Figure 13). The complete blood count 5 months after injection with nanotubes, shows no significant differences between control and nanotube-injected mice (Figure 14-15). Overall, the results suggest minimal effects of nanotubes on any conditions measurable by CBC.

**Figure 11.**
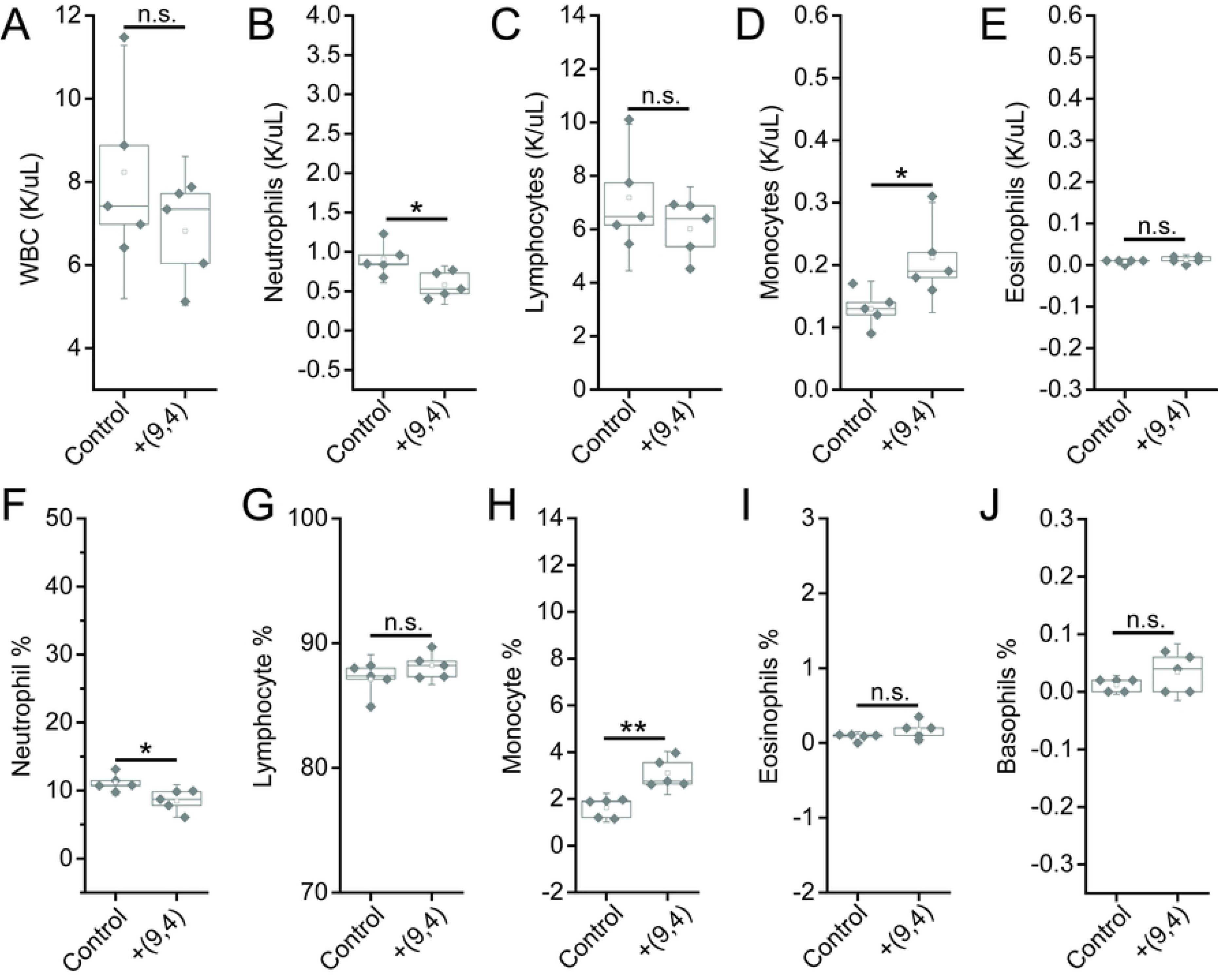
Measurements of blood inflammatory markers in mice 24 hours after injection of nanotubes. Samples were measured 24 hours after injection of PBS (control) or ssCTTC_3_TTC-(9,4) DNA-nanotube complexes. **A)** White blood cell (WBC) concentration in mouse blood. **B)** Neutrophil concentration in mouse blood. **C)** Lymphocyte concentration in mouse blood. **D)** Monocyte concentration in mouse blood. **E)** Eosinophil concentrations in mouse blood. **F)** Neutrophil percentage in mouse blood. **G)** Lymphocyte percentage in mouse blood. **H)** Monocyte percentage in mouse blood. **I)** Eosinophil percentage in mouse blood. **J)** Basophil percentage in mouse blood. * = p < .05 as determined with a Student’s two way t-test. N=5 mice per group.

**Figure 12.**
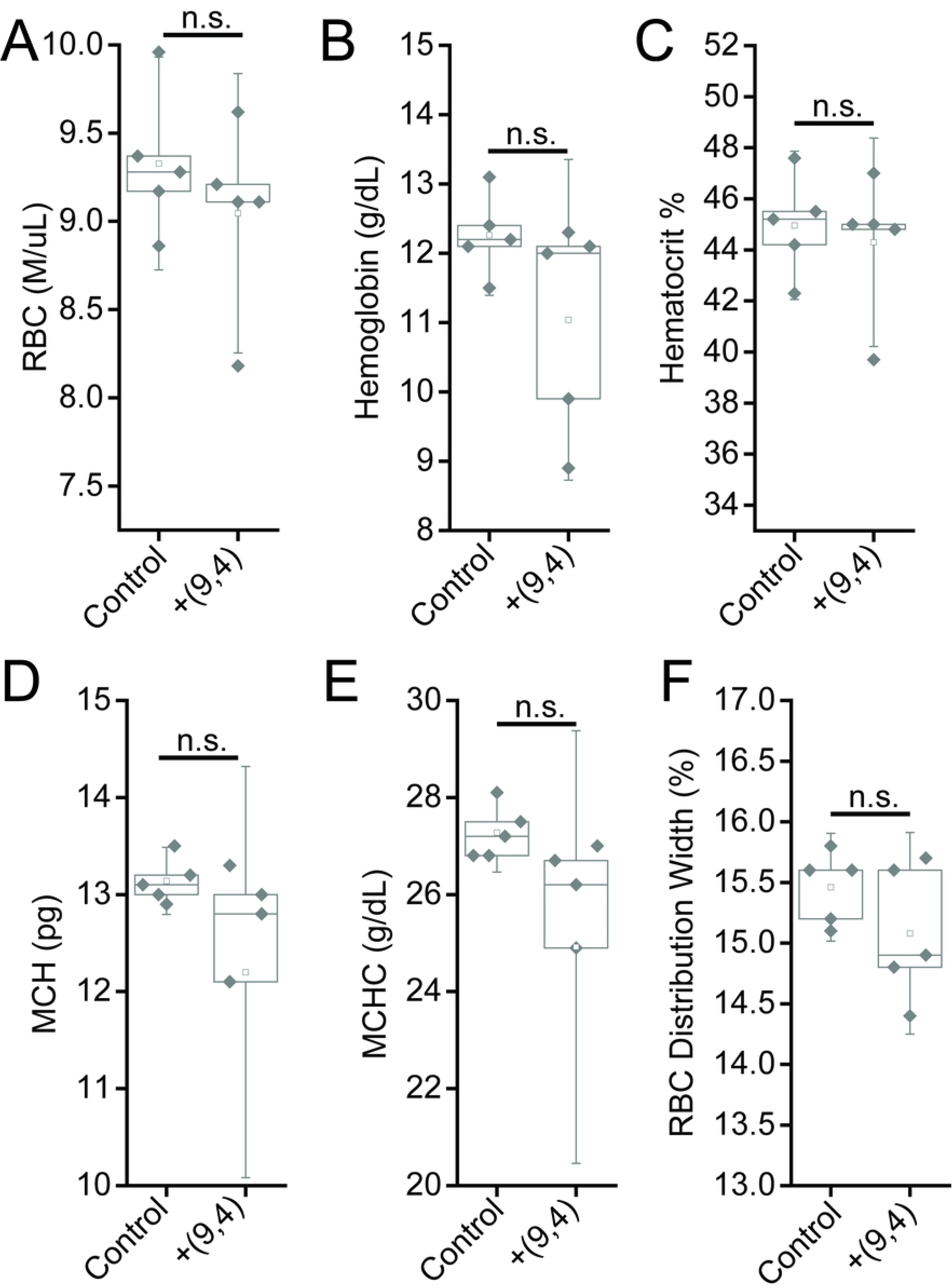
Measurements of blood oxygenation markers in mice 24 hours after injection of nanotubes. Samples were measured 24 hours after injection of PBS (control) or ssCTTC_3_TTC-(9,4) DNA-nanotube complexes. **A)** Red blood cell (RBC) concentration in mouse blood. **B)** Hemoglobin concentration in mouse blood. **C)** Hematocrit percentage in mouse blood. **D)** Mean corpuscular hemoglobin quantity in mouse blood. **E)** Mean corpuscular hemoglobin concentration in mouse blood. **F)** Red blood cell (RBC) distribution width. Statistical significance was determined with a Student’s two way t-test. N=5 mice per group.

**Figure 13.**
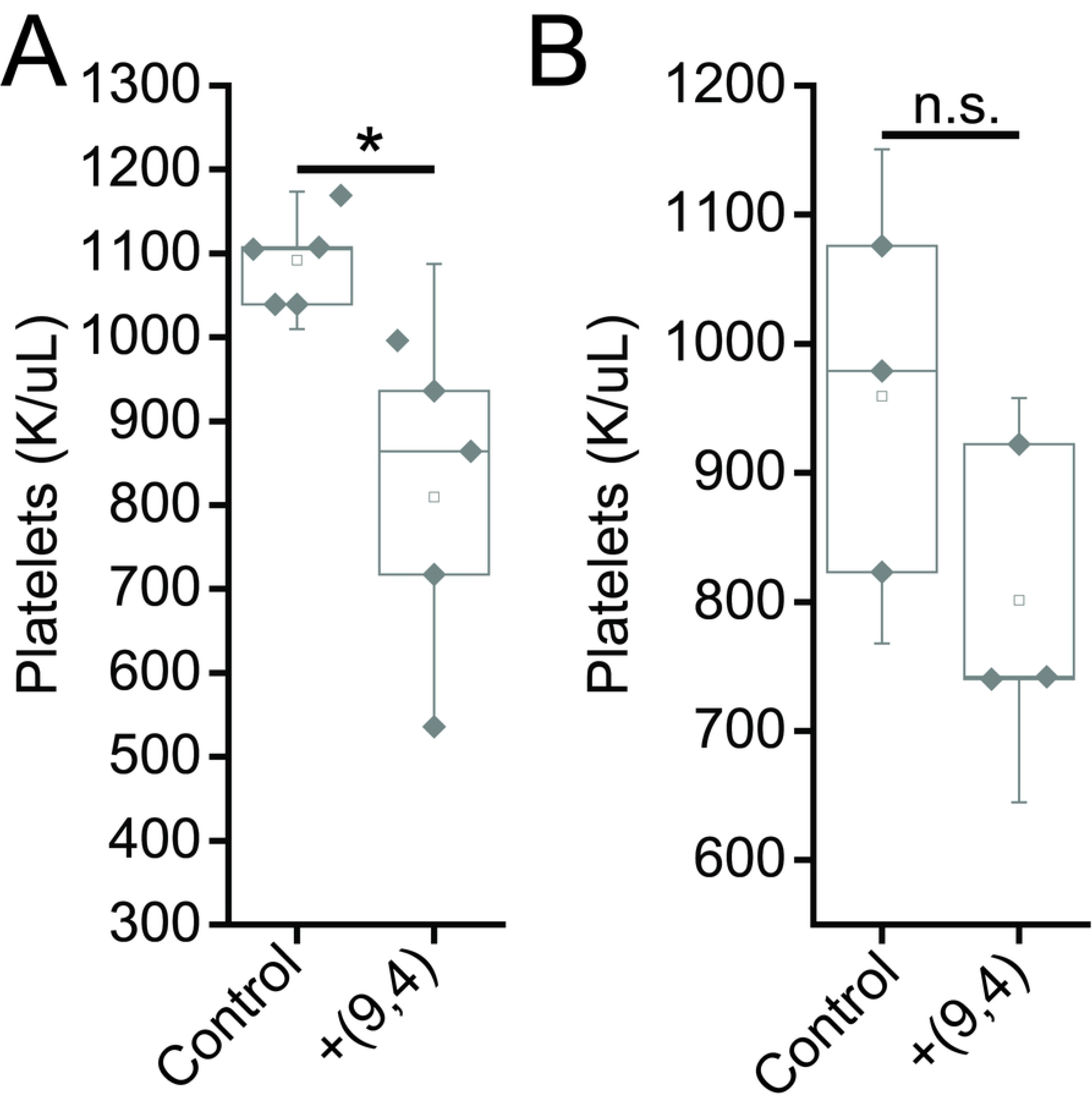
Platelet counts in mice 24 hours and 5 months after injection of nanotubes. Platelet counts were measured 24 hours (A) and 5 months (B) after injection of PBS (control) and ssCTTC_3_TTC-(9,4) complexes.

**Figure 14.**
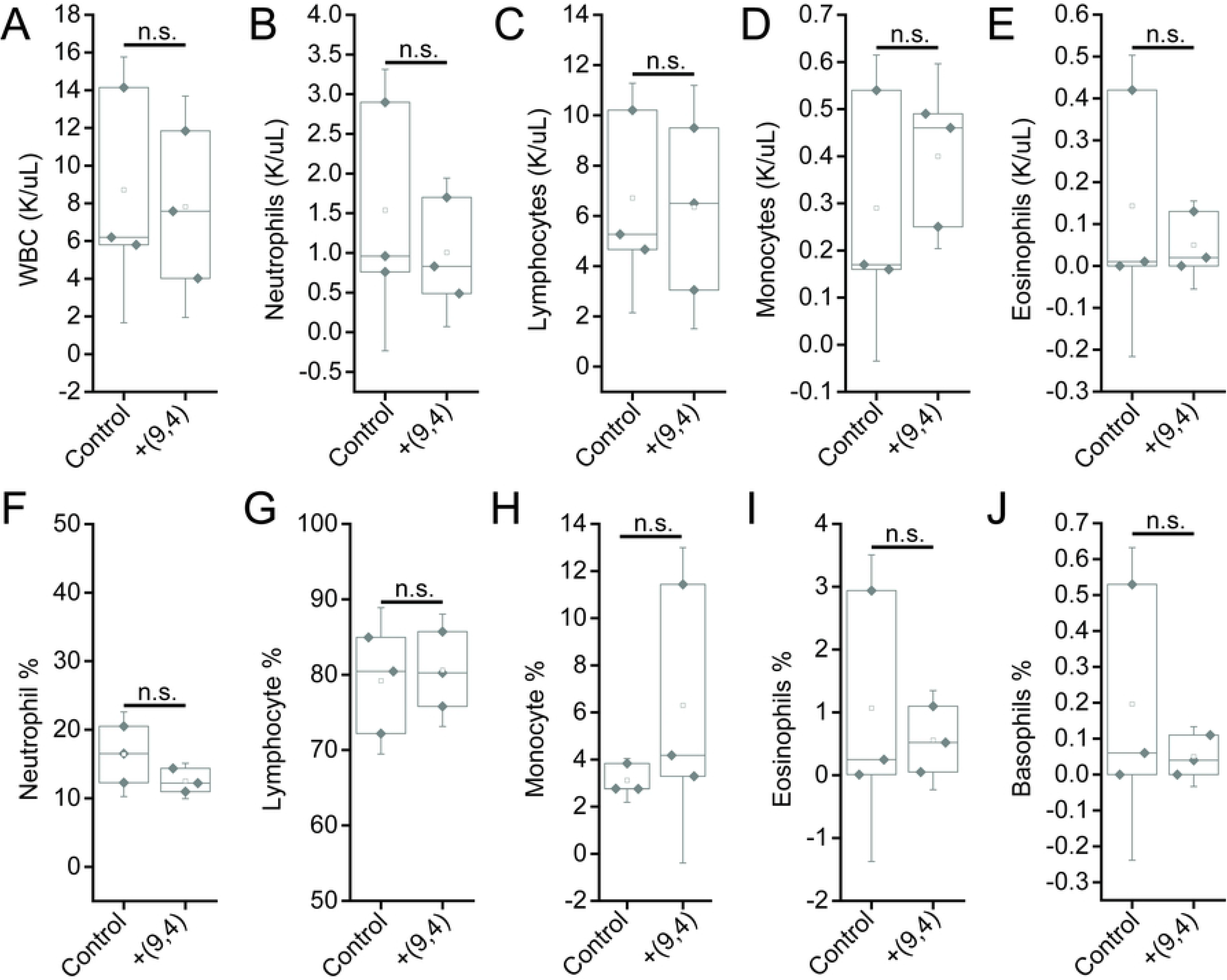
Measurements of blood inflammatory markers in mice 5 months after injection of nanotubes. Samples were measured 5 months after injection of PBS (control) or ssCTTC_3_TTC-(9,4) DNA-nanotube complexes. **A)** White blood cell (WBC) concentration in mouse blood. **B)** Neutrophil concentration in mouse blood. **C)** Lymphocyte concentration in mouse blood. **D)** Monocyte concentration in mouse blood. **E)** Eosinophil concentrations in mouse blood. **F)** Neutrophil percentage in mouse blood. **G)** Lymphocyte percentage in mouse blood. **H)** Monocyte percentage in mouse blood. **I)** Eosinophil percentage in mouse blood. **J)** Basophil percentage in mouse blood. Statistical significance was determined with a Student’s two way t-test. N=3 mice per group.

**Figure 15:**
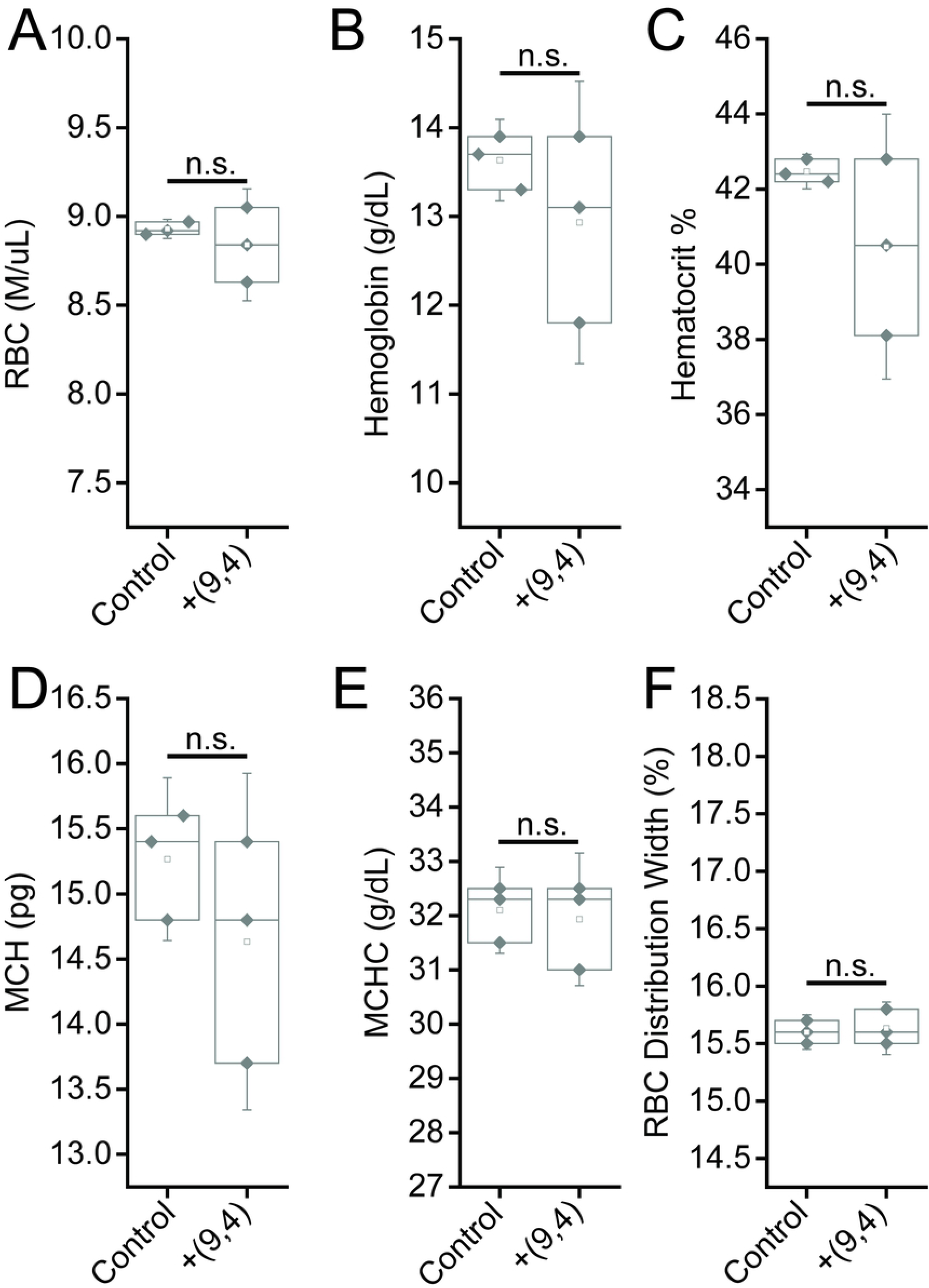
Measurement of blood oxygenation markers in mice 5 months after injection of nanotubes. Samples were measured 5 months after injection of PBS (control) or ssCTTC_3_TTC-(9,4) DNA-nanotube complexes. **A)** Red blood cell (RBC) concentration in mouse blood. **B)** Hemoglobin concentration in mouse blood. **C)** Hematocrit percentage in mouse blood. **D)** Mean corpuscular hemoglobin quantity in mouse blood. **E)** Mean corpuscular hemoglobin concentration in mouse blood. **F)** Red blood cell (RBC) distribution width. Statistical significance was determined with a Student’s two way t-test. N=3 mice per group.

## 4. Conclusion

In this work, we investigated the short and long term biodistribution and biocompatibility of a purified DNA-encapsulated single-walled carbon nanotube complex consisting of an individual nanotube chirality, administered intravenously. Bulk biodistribution measurements in mice found that, consistent with previous studies on similar complexes, the nanotubes localized predominantly to the liver. Using near-infrared hyperspectral microscopy to image single nanotube complexes, nanotube complexes were found in other organs such as the kidney, spleen, hearts, and lungs and persisted in some organs at for up to 5 months. The results showed that this reporter was highly biocompatible overall, although future studies are warranted to more carefully assess long-term impact on organ function. Tissue histology and mouse weight measurements showed no differences upon administration of nanotubes. Measurements of serum biomarkers, including complete blood count, renal biomarkers, and hepatic markers showed negligible changes by the presence of carbon nanotubes < 4 months, and minor changes in renal markers at 5 months. These results indicate that carbon nanotubes, used in preclinical studies under the preparation conditions and concentrations herein, do not cause any appreciable signs of toxicities at time points less than 4 months. The work suggests that these materials are unlikely to cause significant problems in applications such as preclinical research, drug screening, and drug development. However, the long-term persistence of nanotubes in tissues suggests that additional assessments are warranted to assess the potential for use in humans. In general, these studies also suggest, in light of previous works, that carbon nanomaterial biodistribution and biocompatibility are specific to carbon nanotube type, purity, functionalization, and route of administration. This issue has implications pertinent to the wider perception and applicability of nanomaterials.

## Acknowledgements

We would like to thank Y. Shamay, R. Williams, H. Baker, C. Cupo, M. Kim and J. Budhathoki-Uprety for helpful discussions. P.Jena for MATLAB code. We also acknowledge the Molecular Cytology Core Facility at Memorial Sloan Kettering Cancer Center and the Electron Microscopy & Histology Core Facility at Weill Cornell Medicine.

## Supplementary Figure

**Figure S1:**
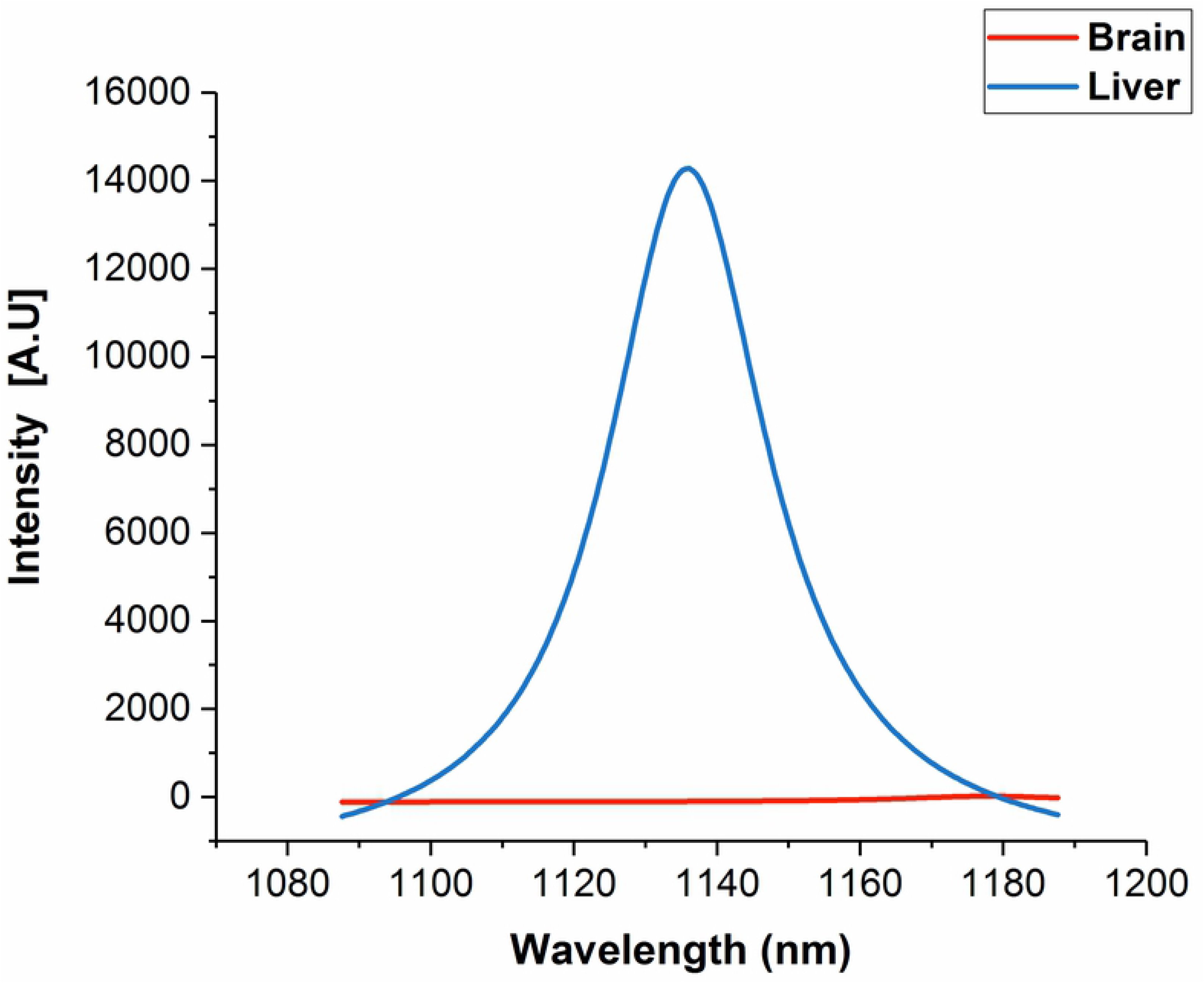
Example near-infrared emission spectra of DNA-SWCNT complexes measured *in vivo* 24 h following intravenous injection into a mouse, in brain and liver tissues.

